# ZmPILS6 is an auxin efflux carrier required for maize root morphogenesis

**DOI:** 10.1101/2023.08.01.551510

**Authors:** Craig L. Cowling, Arielle L. Homayouni, Jodi B. Callwood, Maxwell R. McReynolds, Jasper Khor, Haiyan Ke, Melissa A. Draves, Katayoon Dehesh, Justin W. Walley, Lucia C. Strader, Dior R. Kelley

## Abstract

Plant root systems play a pivotal role in plant physiology and exhibit diverse phenotypic traits. Understanding the genetic mechanisms governing root growth and development in model plants like maize is crucial for enhancing crop resilience to drought and nutrient limitations. This study focused on identifying and characterizing ZmPILS6, an annotated auxin efflux carrier, as a key regulator of various crown root traits in maize. ZmPILS6-modified roots displayed reduced network area and suppressed lateral root formation, desirable traits during drought and low phosphate conditions. The research revealed that ZmPILS6 localizes to the endoplasmic reticulum and plays a vital role in controlling the spatial distribution of indole-3-acetic acid (IAA or “auxin”) in primary roots. The study also demonstrated that ZmPILS6 can actively efflux IAA when expressed in yeast. Furthermore, the loss of ZmPILS6 resulted in significant proteome remodeling in maize roots, particularly affecting hormone signaling pathways. To identify potential interacting partners of ZmPILS6, a weighted gene co-expression analysis (WGNA) was performed. Altogether, this research contributes to the growing knowledge of essential genetic determinants governing maize root morphogenesis, which is crucial for guiding agricultural improvement strategies.

**Significance Statement:** Crop yield and stress resilience are significantly influenced by crown root architecture. A reverse genetic screen aimed at identifying novel regulators of maize root morphogenesis led to the discovery of ZmPILS6, an auxin efflux carrier. The loss of ZmPILS6 negatively impacts numerous root traits that are linked to plant physiology and function. Proteomic characterization of *pils6-1* roots revealed that this evolutionarily conserved transporter is required for the proper expression of numerous phytohormone pathways, including abscisic acid, gibberellins, and jasmonic acid. Notably, ZmPILS6 appears to have a contrasting role in regulating root morphogenesis compared to its Arabidopsis ortholog, PILS6. This finding emphasizes the need for functional characterization of candidate genes directly within key crops of interest, which cannot always be correctly inferred from other model plants.

## Main Text

Plant roots are an important organ for anchorage and uptake of water and nutrients. Root development is influenced by plant hormone transport dynamics to influence growth and differentiation (1–4). In Arabidopsis, an asymmetric distribution of indole-3-acetic acid (IAA, or “auxin”) across the primary root is required for proper cell division and patterning (5–15). Auxin transport is carried out by several evolutionarily conserved proteins, including the plant-specific PIN-FORMED (PIN) family, PIN-likes (PILS), ATP-binding cassette (ABC) B-type (ABCB) family, AUX1/LAX family (16, 17). In maize, the roles of auxin transport during embryogenesis, leaf development, and inflorescence architecture have been well established (18–22). However, the roles of auxin transport in maize root morphogenesis are not well understood. Orthologs of Arabidopsis PIN1, ZmPIN1 genes, are polarly localized and required for organ formation(20, 23). The four ZmPIN1 genes exhibit tissue-specific expression patterns, suggesting sub-functionalization among the family members (24).

In Arabidopsis, PIN proteins are typically associated with the plasma membrane while PILS and non-canonical PINs are localized to the endoplasmic reticulum(16). These proteins control intra- and intercellular IAA transport and are required for proper root growth and development (25, 26). Loss of *PILS6* in Arabidopsis leads to increased primary root length and elevated expression of *CYCB1;1:GUS* expression, indicating a negative role for PILS in root morphogenesis (26). Although PILS proteins are evolutionarily conserved among land plants, their roles outside of eudicots are yet to be determined (26–28). Several of the nine annotated ZmPILS genes are induced in maize roots in response to abiotic stress (29) and many of the family members exhibit tissue specific expression patterns (30). ZmPILS4 was identified as a major QTL of xylem traits in maize (31), suggesting that PILS proteins may have shared and unique roles in root development.

Reverse genetic screens are an effective approach for linking gene to function in eukaryotic systems (32). Here, we leveraged an integrated gene expression atlas (30) in order to identify putative auxin transporters that are required for maize root formation. By focusing on candidate annotated auxin efflux carriers with enriched protein abundance in maize primary roots and utilizing publicly available transposon insertion lines (33–35) this screen identified ZmPILS6 (Zm00001eb149720, also previously called ZmPILS4) as an influencer of root morphogenesis. Based on phylogenetic analysis of Arabidopsis and maize PILS proteins, Zm00001eb149720 is most closely related to AtPILS6 (At5g01990) and is thus designated ZmPILS6. Loss of *ZmPILS6* leads to reduced crown root architecture and diminished lateral root formation. In addition, ZmPILS6 is localized to the endoplasmic reticulum and is required for proper transport of IAA. A proteomic analysis revealed that several decreased proteins in *zmpils6* relative to wild-type W22 are involved in hormone pathways, including auxin. Using these data, a weighted gene co-expression network analysis (WGCNA) was performed to identify candidate proteins that may act with ZmPILS6 to influence maize root development. Altogether this novel work establishes roles for auxin transport in maize root formation and suggests PILS proteins may have opposing functions in monocots and eudicots.

## Results and Discussion

### Loss of ZmPILS6 negatively impacts crown root architecture

To identify candidate genes for a reverse genetics screen on auxin transporters in maize roots, we utilized available quantitative proteomics data for maize (30). A query of the maize expression atlas identified ZmPILS6 (Zm00001eb149720) as an annotated auxin efflux carrier with enriched expression in unpollinated silks, internodes, and primary roots (*SI Appendix*, Fig. S1). ZmPILS6 is the ortholog of Arabidopsis PILS6 based on a phylogenetic analysis of full-length PILS proteins sequences from TAIR and MaizeGDB (*SI Appendix*, Fig. S2). Three Mu transposon alleles were identified for ZmPILS6 from publicly available stocks, which are designated as *pils6-1* (mu1047700), *pils6-2* (mu1090629), and *pils6-3* (mu-ill 221882.6) (Fig. 1A). Both *pils6-1* and *pils6-2* are W22 Uniform Mu lines with transposon insertions in the 5’ UTR, while *pils6-3* is a B73 Mu Illumina line with a transposon insertion in the first exon (Fig. 1A). *ZmPILS6* expression was examined in all three alleles and their respective inbred background using RT-qPCR and found to be significantly reduced in both *pils6-1* and *pils6-3* alleles, but not significantly changed in the *pils6-2* allele (Fig. 1B). Based on this expression data, the *pils6-2* allele was not used further in this study.

**Figure 1.**
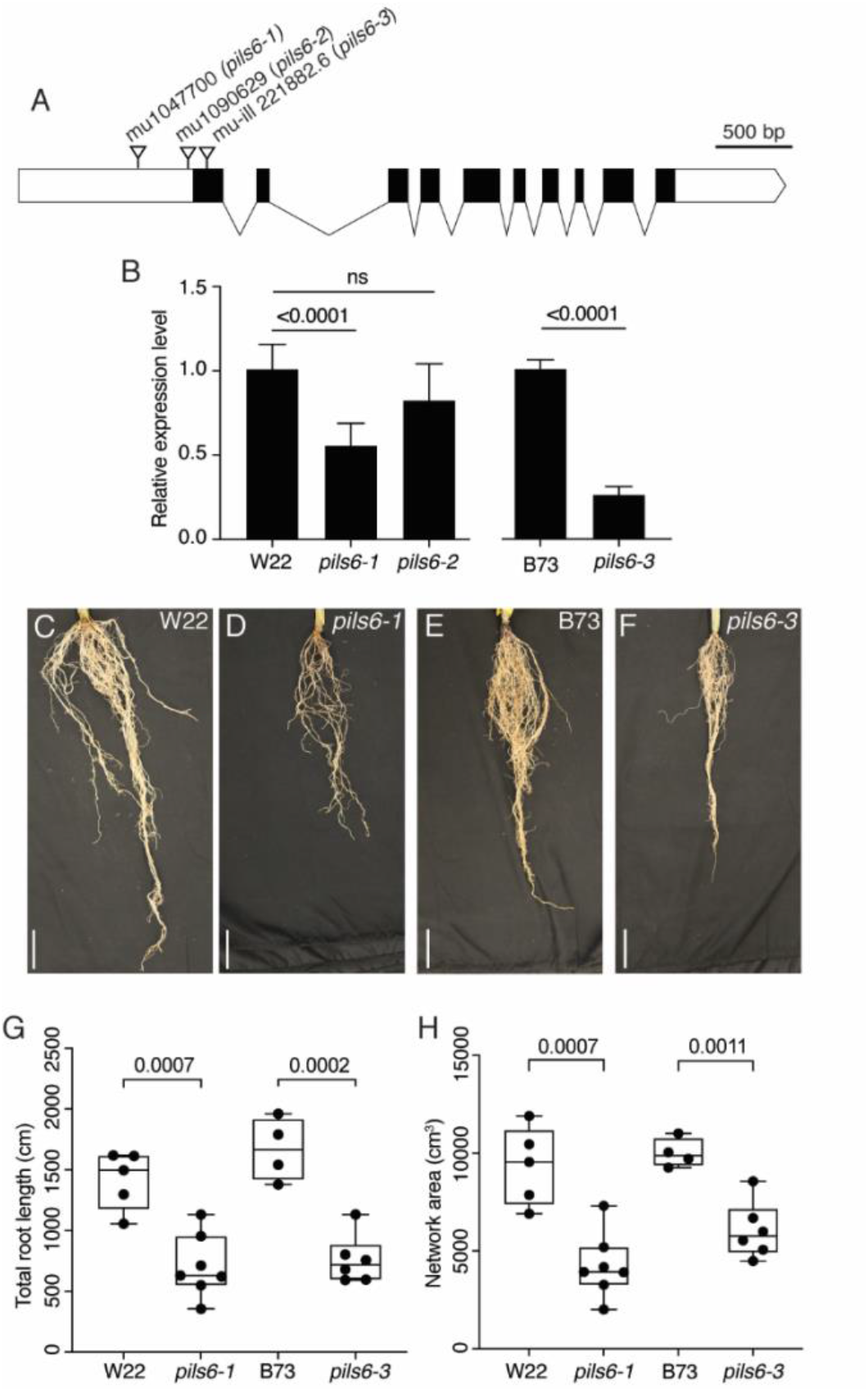
Characterization of *ZmPILS6* loss-of-function alleles. (A) Schematic of the *ZmPILS6* gene (Zm00001eb149720). White boxes indicate untranslated regions (UTR), exons are diagramed in black boxes, and introns are shown as lines. The location of *ZmPILS6* transposon alleles are shown as white inverted triangles. The alleles characterized in this study are designated as *pils6-1* (mu1047700), *pils6-2* (mu1090629), and *pils6-3* (mu-ill 221882.6). (B) *ZmPILS6* expression is reduced in *pils6-1* and *pils6-3* based on RT-qPCR. (C-F) Representative images of crown root phenotypes in W22 (C), *pils6-1* (D), B73 (E), and *pils6-3* (F). Crown root quantification of total root length (G) and network area. Scale bars = 5 cm.

A phenotypic analysis of crown root systems of wild-type W22 (Fig. 1C) and B73 (Fig. 1E) compared to the knock-down *ZmPILS6* alleles determined that root morphogenesis is greatly reduced in both *pils6-1* (Fig. 1D) and *pils6-3* (Fig. 1F). Furthermore, both *pils6-1* and *pils6-3* exhibited a significant reduction in total root length (Fig. 1G) and network area (Fig. 1H) compared to W22 and B73. Loss of *ZmPILS6* also led to a reduction in branch points and fewer root tips (*SI Appendix*, Fig. S3, Dataset S1). These key root characteristics are important traits of agricultural relevance (36) and demonstrate that ZmPILS6 is a key genetic determinant of maize root systems. Specifically, ZmPILS6 is a positive regulator of root morphogenesis in maize. This result contrasts with the known roles of the Arabidopsis PILS6 ortholog, which is a negative regulator of root growth and development (26).

### ZmPILS6 is required for lateral root formation

In Arabidopsis, PILS2 and PILS5 are known to regulate lateral root formation (25). In order to determine if ZmPILS6 plays a role in this same process, lateral root formation was examined in six-day-old roots of *pils6-1* and *pils6-3* using an established Feulgen staining protocol (37). Compared to wild-type W22 (Fig. 2A), the mature zone of *pils6-1* roots exhibited fewer lateral roots (Fig. 2B). This same phenotype was also observed in *pils6-3* (Fig. 2D) compared to B73 (Fig. 2C). Both knock-down alleles had a reduction in lateral root primordia density, which was determined as the number of primordia divided by the length of the primary root (Fig. 2E,F). These results demonstrate that ZmPILS6 is a positive regulator of lateral root development in maize.

**Figure 2.**
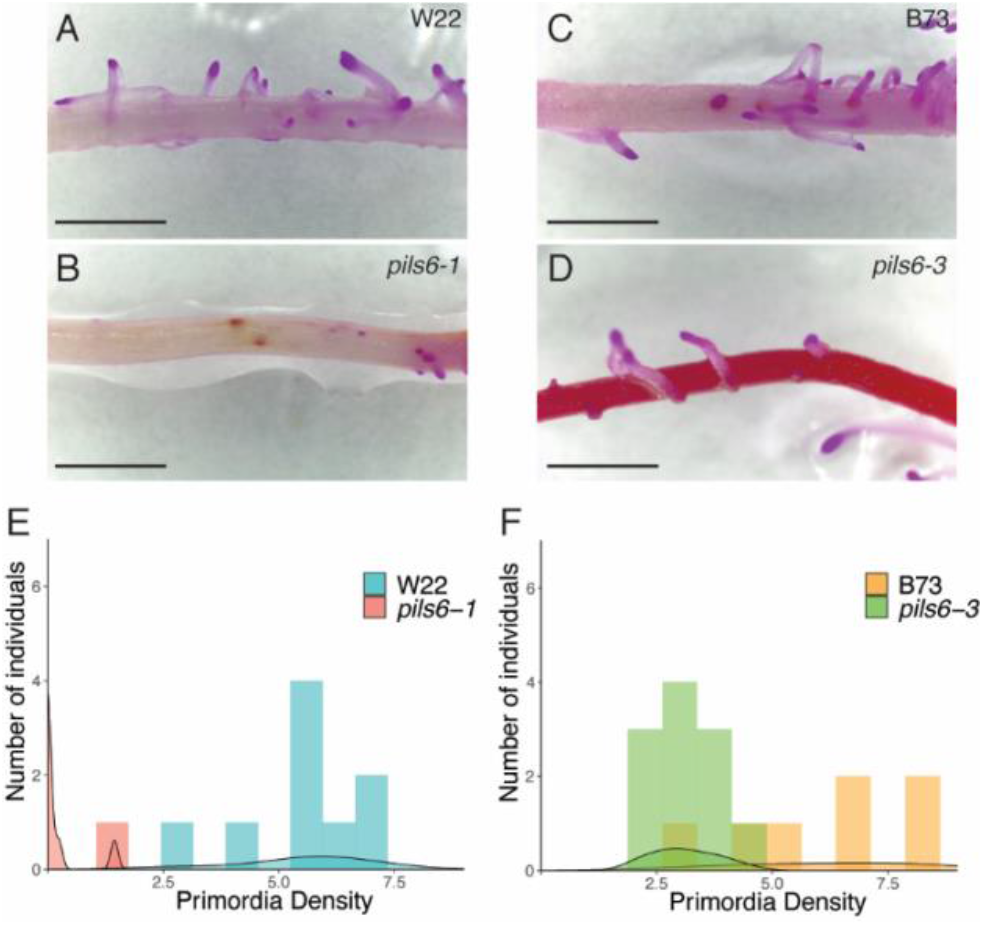
Lateral root formation is reduced in *pils6* mutants. Feulgen stained six-day-old primary roots of W22 (A), *pils6-1* (B), B73 (C), and *pils6-3* (D). Scale bars = 2 mm. (E-F) Histogram visualization of lateral root primordia density in *pils6* alleles compared to their respective inbred controls. Scale bars = 2 mm.

### ZmPILS6 can transport indole-3-acetic acid

Auxin distribution and response is non-uniform across the maize primary root (38–40). To further investigate this pattern endogenous levels of IAA were quantified using LC-MS on five-day-old primary roots hand dissected into four regions: the meristematic zone (MZ), elongation zone (EZ), cortex, and stele (which were obtained from the differentiation zone) (Fig. 3A, Dataset S2). Endogenous IAA levels were highest in the stele and meristematic zone, while the elongation zone and cortex contained comparable amounts of IAA (Fig. 3A). This asymmetric pattern could be established due to tissue specific auxin biosynthesis, transport and/or metabolism. To determine if ZmPILS6 can contribute to IAA transport in primary roots, a modified radiolabeled ^3^H-IAA assay was performed using wild-type and *pils6-1* dissected roots. In the absence of ZmPILS6, ^3^H-IAA hyperaccumulates the meristematic and elongation zones, which is consistent with a defect in efflux of this molecule (Fig. 3B). ZmPILS6 is localized to the endoplasmic reticulum in *Nicotiana benthimiana* cells (Fig. 3C), which is consistent with the known subcellular localization of Arabidopsis PILS orthologs. To establish if ZmPILS6 is capable of directly transporting ^3^H-IAA, full-length ZmPILS6 protein was expressed in yeast. In these assays we observed a significant decrease in ^3^H-IAA accumulation in yeast cells expressing ZmPILS6 compared to the empty vector control, which is consistent with ZmPILS6 efflux activity in yeast (Fig. 3D). Altogether these data suggest that ZmPILS6 is capable of directly effluxing IAA from certain tissues, likely across the ER membrane. Furthermore, the spatial dynamics of IAA efflux in maize primary roots is in part conferred by ZmPILS6. In *pils6-1* roots, auxin efflux is specifically altered within the meristematic and elongation zones of the primary root, but normal within the maturation zone.

**Figure 3.**
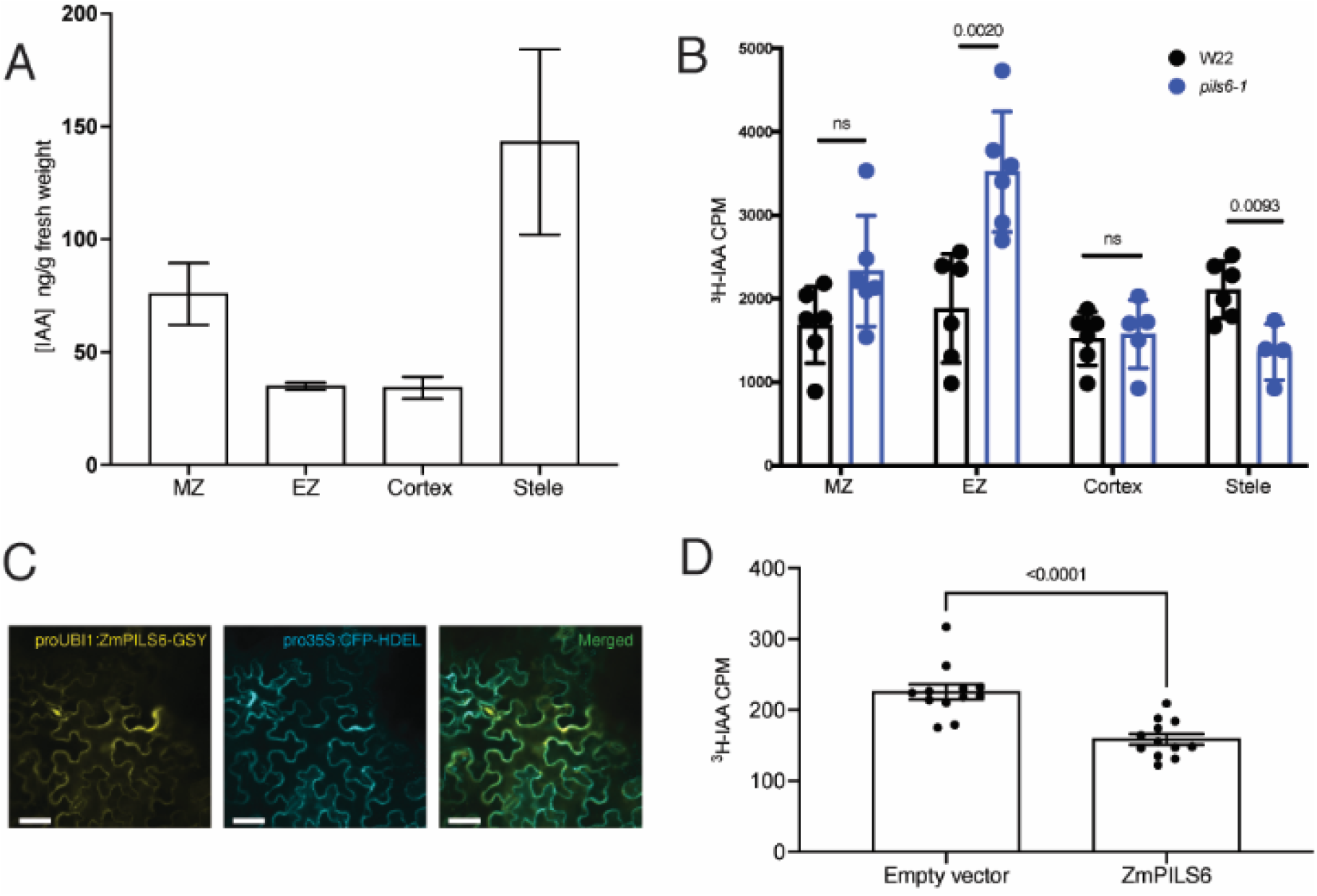
ZmPILS6 is an ER localized auxin efflux protein. (A) Indole-3-acetic acid (IAA) levels in dissected regions of five-day-old primary roots: meristematic zone (MZ), elongation zone (EZ), cortex, and stele. (B) Levels of ^3^H-IAA in dissected regions of primary W22 and *pils6-1* roots. *P* values shown from ANOVA; ns = not significant. (C) Transient expression of *pUBI1:PILS6-GS*^*yellow*^ (yellow fluorescence) and a marker for the endoplasmic reticulum (ER), *p35S:CFP-HDEL* (blue fluorescence), in *Nicotiana benthamiana* epidermal cells. Overlay of both channels is shown in green (“merged”). (D) Levels of ^3^H-IAA in yeast expressing an empty vector or ZmPILS6. Scale bars = 50 μm.

### Loss of *ZmPILS6* impacts auxin responsive proteome remodeling

Auxin signaling can rapidly and extensively impact proteome dynamics in numerous angiosperm organs (41–44). Proteomic profiling of *pils6-1* and W22 roots was performed in the presence and absence of auxin treatment (10 μM IAA) to identify protein expression patterns in the mutant (Dataset S3). In W22, 178 proteins exhibited altered abundance in response to IAA treatment. In comparison, auxin responsiveness in *pils6-1* is reduced with only 9 auxin-responsive proteins. Overall, greater changes in proteome composition were observed between *pils6-1* and W22 in the absence of auxin, whereby thousands of proteins were increased or decreased compared to W22 (Fig. 4A). Notably, a comparison of the auxin-responsive proteins between the genotypes uncovered a greater influence on the proteome. Specifically, 1493 proteins are constitutively increased in *pils6-1* compared to W22 following auxin treatment, while 1174 proteins remained repressed in *pils6-1* roots in the presence of auxin (Fig. 4B). In addition, discordant protein abundance changes were observed between auxin-responsive proteins in *pils6-1* and W22 (Fig. 4B). For example, 11 proteins which are auxin-induced in W22 are auxin-repressed in *pils6-1*.

**Figure 4.**
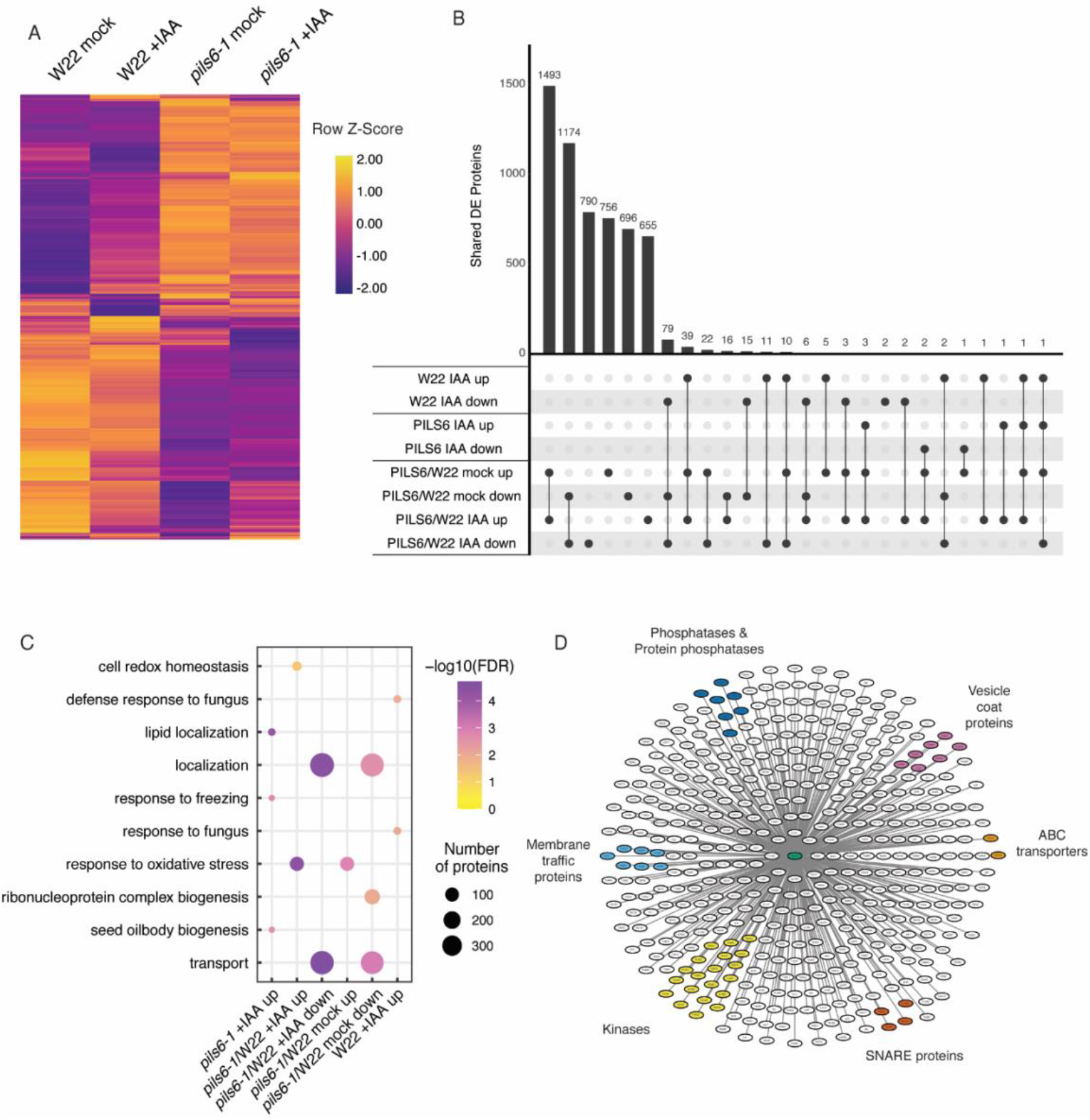
Proteome changes in the absence of ZmPILS6. (A) Hierarchical clustering of differentially expressed (DE) proteins in W22 and *pils6-1*, both in the presence and absence of auxin. (B) Upset plot of DE proteins indicates extent of overlap between DE proteins across genotypes and treatments. (C) Gene ontology enrichment of DE proteins in W22 and *pils6-1*. (D) A ZmPILS6 co-expression protein network reconstructed from weighted gene co-expression network analysis. Nodes are colored according to PANTHER protein class, whereby ZmPILS6 (central green node) is predicted to interact with 414 co-expressed proteins, including vesicle coat proteins (reddish purple), ABC transporters (orange), SNARE proteins (vermillion), kinases (yellow), membrane traffic proteins (sky blue), and phosphatases (blue).

Proteins involved in transport and localization are overrepresented among DE proteins in *pils6-1* roots (Fig. 4C, Dataset S4). In addition to the enriched GO terms, several hormone related proteins were impacted in *pils6-1* including auxin biosynthesis and transport, gibberellic acid response, abscisic acid responsive transcription factors, brassinosteroid related kinases, and jasmonic acid proteins associated with plant development (Dataset S3). Using these proteomics data, a weighted gene co-expression network analysis was performed to predict and identify proteins that may be linked to ZmPILS6, which identified 19 modules (*SI Appendix*, Fig. S4). The ‘greenyellow’ module contains 1299 highly correlated proteins, including ZmPILS6 and its paralog ZmPILS2 (Dataset S5). In addition, there were 415 proteins that were highly correlated to ZmPILS6 based on our proteomics data (Fig. 4D). These include vesicle coat proteins, SNAP receptors (SNARE) proteins, kinases, phosphatases, and ABC transporters, which are color-coded in the network (Fig. 4D, Dataset S6). The regulation of polar auxin transport by AGC kinases and several distinct LRR-RLKs (45–47) and ZmPILS6 is known to be phosphorylated in vivo (30). In addition, interactions between Arabidopsis PINs and ABCBs influence protein complex stability and function (48, 49). The identification of SNARE and vesicle coat proteins in the ZmPILS6 network is consistent with several reports on polarization of PIN proteins by SNAREs (50–52), and suggests that ZmPILS6 may also require SNAREs for proper subcellular localization.

### Concluding Remarks

PILS proteins are evolutionarily conserved among land plants (26, 53). Most land plants contain several PILS orthologs, while sister algal groups contain only 1-2 annotated PILS genes (26). This radiation of the PILS gene family during land plant evolution may have been accompanied by neo- or sub-functionalization, which will require future investigation into the individual roles of PILS proteins across different land plants. The ER-localization and ability to contribute to auxin transport appears to be conserved between maize and Arabidopsis, suggesting evolutionarily conserved properties of these PILS. However, while this study supports a role for Clade II (PILS2/6 clade) proteins in regulating root formation, there are apparent diverged functions between Arabidopsis (a core eudicot) and maize (a monocot). PILS proteins in Arabidopsis are negative regulators of root growth but ZmPILS6 is a positive driver of root system architecture. Future investigation into PILS interacting proteins and their associated downstream processes will be required to determine the mechanisms underpinning these functional differences *in planta*. In addition, the identification of ZmPILS6 several co-expressed kinases, ABC transporters, and SNAREs will enable continued studies into protein complexes that influence auxin transport in maize.

## Materials and Methods

### Plant materials

UniformMu and mu-illumina seed stocks were obtained from the Maize Genetics Cooperation Stock Center and genotyped using gene specific primers for *ZmPILS6* (*Zm00001eb149720*) (*SI appendix*, Table S1). Alleles were designated as *pils6-1* (mu1047700), *pils6-2* (mu1090629), and *pils6-3* (mu-ill 221882.6). Maize kernels were surface sterilized and planted as previously described (54). Auxin response assays were performed as previously described (54) using 20-40 biological replicates per genotype and treatment.

### Crown root phenotyping

Maize seedlings were grown in greenhouse conditions to the V7 stage in 1-gallon pots with soil. Seedlings were removed from the pots and root systems were rinsed with tap water to remove all soil. After drying at room temperature, crown root systems were photographed using a Cannon DSLR camera. Crown root images were analyzed using Rhizhovision Explorer (36). Statistical analysis of crown root traits was performed using a t-test in R using the t.test function.

### Lateral root primordia staining

Kernels were sterilized and planted as described. At six days after germination (DAG), primary roots were incubated in Feulgen stain as previously described (37). For each genotype,7-11 biological replicates were stained per genotype. Lateral roots and lateral root primordia were counted by eye and statistical analysis was performed using the t.test function in R.

### RNA Extraction

Maize kernels were grown until seedlings were 5 DAG as described above. For each biological replicate, primary root tissue from 5 individual seedlings was harvested and pooled together, flash frozen in liquid nitrogen, and ground to a homogenous fine powder using a mortar and pestle. Total RNA was extracted from # of biological replicates per genotype. Total RNA was extracted as previously described (40). The purified total RNA was used to synthesize cDNA using LunaScript® RT SuperMix Kit (New England Biolabs, catalog number E3010).

### RT-qPCR Assay

15 ng of cDNA was used per reaction to amplify ZmPILS6 from root tissues. Brown midrib4/FPGS (Zm00001eb404110) was used as a control reference gene. Data analysis was done using the delta delta Ct method as previously described (55).

### Cloning

All plasmids were verified using restriction enzyme digestions and whole-plasmid sequencing. Full length *ZmPILS6* was cloned into pENTR/D-TOPO using primers in *SI appendix*, Table S1. The resulting entry clone driven by the promoter UBI1, and a C-terminal GS^yellow^ fluorescent tag were transferred to the Gateway-compatible destination vector, pBb7m34GW (56) to create plasmid pLD19. pLD19 was transformed into One Shot TOP10 E coli and screened on LB plates containing spectinomycin (100 ug/mL). pML1 was recombined into pAG426GPD-ccdb using LR Clonase to create pCC13.

### Transient expression and confocal microscopy

*Agrobacterium tumefaciens* strain GV3101 was transformed with pLD19 via the freeze-thaw method. Bacteria was grown in liquid LB medium containing spectinomycin (100 ug/mL) or kanamycin (100 ug/mL) overnight at 30°C. Cells were resuspended in MES Buffer (10mM MES, 10mM MgCl_2_, PH 5.7) and cell density was adjusted to an OD600 of 0.3. Plasmid containing bacteria was mixed with P19 bacteria in a 1:1 ratio with a final OD600 of 0.4 (57). Bacteria was infiltrated into 4-week-old *Nicotiana benthiamiana* plants using a needleless syringe. Transient expression was observed 3 days post inoculation using a Zeiss LSM 700 confocal microscope. CFP fluorescent signals were captured at 485 nm after excitation of 405 nm. GS^yellow^ signals were captured between 525-575 nm after excitation of 514 nm. Transient assays were repeated in triplicate.

### Auxin transport assays in yeast

Transport assays of radiolabeled indole-3-acetic acid (^3^H-IAA) were performed as previously described (58). Briefly, pCC13 was transformed into *Saccharomyces cerevisiae* strain JBY575. Single yeast colonies were grown in liquid -URA selective media overnight at 30°C. Yeast were resuspended in 0.1M MES (pH 4.6) with 2% dextrose to an OD600 of 5. 100 μL of resuspended cells were added to a 1.5mL Eppendorf tube and added 100 μL of 50 nM ^3^H-IAA and incubated at RT for 30 minutes. Yeast cells were centrifuged for 15 seconds before removing the supernatant and washing 3 times with MES solution. Cells were resuspended in 100 uL of MES buffer and ^3^H-IAA was measured using a scintillation counter. For each plasmid expressing yeast, 12 biological replicates were statistically analyzed using a t test.

### *In vivo* auxin transport assays

Auxin accumulation assays were performed as previously described (59). Maize seedlings were grown for 6 days and dissected into four regions as previously described (40, 60), designated the meristematic zone (“MZ”), elongation zone (“EZ”), cortical parenchyma and epidermis (“cortex)”, and vasculature (“stele”). Sections were equilibrated in 40 μL of uptake buffer (20 mM MES, 10 mM sucrose, and 0.5 mM CaSO4, pH 5.6) for 30 minutes at room temperature (RT). 40 μL of 50 nM ^3^H-IAA was added to each sample and incubated at RT for 1 hour. Samples were rinsed 3 times with 80 μL of uptake buffer. ^3^H-IAA in tissue samples was measured using a scintillation counter.

### Indole-3-acetic acid measurements

Five-day-old B73 primary roots were hand dissected into meristematic zone (MZ), elongation zone (EZ), cortex, and stele and pooled to reach 200 mg per root region and replicate. The tissue was flash frozen and ground into a uniform powder using a mortar and pestle under liquid nitrogen. Extraction and measurements were done as previously described using isopropanol:water:HCl (2:1:2) as the extraction buffer (61) with 50 mg of ground root tissue per replicate. Extracts were analyzed via liquid-chromatography mass spectrometry (LC-MS) on a Thermo Dioned Ultimate 3000 + Q-Exactive Focus run with an internal indole-3-acetic acid (IAA) sample. For each sample, 5-6 biological replicates were analyzed (Dataset S2).

### Proteomics

#### Protein Extraction and Digestion

The proteomics experiments were carried out based on established methods (62). Protein was extracted and digested into peptides with trypsin and Lys-C using the phenol-FASP method as previously detailed (63, 64). Resulting peptides were desalted using 50 mg Sep-Pak C18 cartridges (Waters), dried using a vacuum centrifuge (Thermo), and resuspended in 0.1% formic acid. Peptide amount was quantified using the Pierce BCA Protein assay kit.

#### Tandem Mass Tag (TMT) Labeling

TMTpro™ 16plex labeling reagents (ThermoFisher, Lot VB294909) were used to label at a TMT:peptide ratio of 0.2:1 as described in (Song et al., 2020). After 2 hours incubation at room temperature the labeling reaction was quenched with hydroxylamine. Next, the 16 samples were mixed together stored at -80ºC.

#### LC-MS/MS

An Agilent 1260 quaternary HPLC was used to deliver a flow rate of ∼600 nL min-1 via a splitter. All columns were packed in house using a Next Advance pressure cell, and the nanospray tips were fabricated using a fused silica capillary that was pulled to a sharp tip using a laser puller (Sutter P-2000). 10 μg of TMT-labeled peptides were loaded onto 10 cm capillary columns packed with 5 μM Zorbax SB-C18 (Agilent), which was connected using a zero dead volume 1 μm filter (Upchurch, M548) to a 5 cm long strong cation exchange (SCX) column packed with 5 μm PolySulfoethyl (PolyLC). The SCX column was then connected to a 20 cm nanospray tip packed with 2.5 μM C18 (Waters). The 3 sections were joined and mounted on a Nanospray Flex ion source (Thermo) for on-line nested peptide elution. A new set of columns was used for every sample. Peptides were eluted from the loading column onto the SCX column using a 0 to 80% acetonitrile gradient over 60 minutes. Peptides were then fractionated from the SCX column using a series of 1 8 salt steps (ammonium acetate) for the non-modified proteome and phosphoproteome analysis, respectively. For these analyses, buffers A (99.9% H_2_O, 0.1% formic acid), B (99.9% ACN, 0.1% formic acid), C (100 mM ammonium acetate, 2% formic acid), and D (2 M ammonium acetate, 2% formic acid) were utilized. For each salt step, a 150-minute gradient program comprised of a 0–5 minute increase to the specified ammonium acetate concentration, 5–10 minutes hold, 10–14 minutes at 100% buffer A, 15–100 minutes 15–30% buffer B, 100-121 minutes 30–45% buffer B, 120–140 minutes 45–80% buffer B, 140–144 minutes 80% buffer B, and 145–150 minutes buffer A was employed. minutes buffer A was employed.

Eluted peptides were analyzed using a Thermo Scientific Q-Exactive Plus high-resolution quadrupole Orbitrap mass spectrometer, which was directly coupled to the HPLC. Data dependent acquisition was obtained using Xcalibur 4.0 software in positive ion mode with a spray voltage of 2.20 kV and a capillary temperature of 275 °C and an RF of 60. MS1 spectra were measured at a resolution of 70,000, an automatic gain control (AGC) of 3e6 with a maximum ion time of 50 ms and a mass range of 350-1500 m/z. Up to 15 MS2 were triggered at a resolution of 35,000 with a fixed first mass of 110 m/z. An AGC of 2e5 with a maximum ion time of 120 ms, an isolation window of 1.3 m/z, and a normalized collision energy of 31. Charge exclusion was set to unassigned, 1, 5–8, and >8. MS1 that triggered MS2 scans were dynamically excluded for 30 s.

#### Proteomics Data Analysis

The raw spectra were analyzed using MaxQuant version 1.6.14.0 (65). Spectra were searched using the Andromeda search engine in MaxQuant (66) against the *Zea mays* B73 proteome file entitled “Zea_mays.AGPv4.pep.all” and was complemented with reverse decoy sequences and common contaminants by MaxQuant. Carbamidomethyl cysteine was set as a fixed modification while methionine oxidation and protein N-terminal acetylation were set as variable modifications. The sample type was set to “Reporter Ion MS2” with “16plex TMT selected for both lysine and N-termini”. Digestion parameters were set to “specific” and “Trypsin/P;LysC”. Up to two missed cleavages were allowed. A false discovery rate, calculated in MaxQuant using a target-decoy strategy (67), less than 0.01 at both the peptide spectral match and protein identification level was required. The ‘second peptide’ option identify co-fragmented peptides was not used. The match between runs feature of MaxQuant was not utilized.

Statistical analysis on the MaxQuant output was performed using the TMT-NEAT Analysis Pipeline (68). Differential expression (DE) was determined using a q-value □0.05 (Dataset S3). Upset plots were generated in R as previously described (40). Proteins with altered abundance were hierarchically clustered based on their protein abundance under mock or IAA treatments and plotted as heatmaps using the Morpheus software from the Broad Institute.

### Gene ontology analysis

Gene ontology (GO) enrichment analysis was performed at PANTHER using the *Zea mays* B73 inbred reference genome as previously described (40). The DE proteins were tested for statistical over-representation of GO terms using Fisher’s exact test and an FDR correction using the Benjamini-Hochberg method with a cutoff of 0.05 (Dataset S4). Significant GO Biological Process terms were plotted against the genotype/treatments using a multidimensional dot plot in R (69).

### Weighted Gene Co-expression Network Analysis (WGCNA)

A ZmPILS6 co-expression network was created using protein expression from mock and IAA treated samples. 12,265 globally detected protein abundances (Dataset S3) were used as input to the signed WGCNA network construction using the WGCNA v1.70-3 package in R (70). In WGCNA networks, power was set to 9, minModuleSize was set to 100, and initial clusters were merged on eigengenes (Dataset S5). The mergeCutHeight value was set to 0.20 across all networks. Visualization of the ZmPILS6 co-expression network (Dataset S6) was visualized in Cytocape version 3.9.1 using the organic layout.

### Phylogenetic analysis

Sequences for Arabidopsis PILS proteins and maize PILS proteins were obtained from PLAZA (71) to construct the phylogenetic tree.

## Data availability

Raw proteomics data have been deposited on MassIVE with accession number MSV000092438.

## Acknowledgments

We wish to thank Erik Vollbrecht and Lander Geadelmann for sharing *pils6-3* and B73 kernels.

## Notes

### Competing Interest Statement

The authors have declared no competing interest.

