## Supplementary figures and images for "ZmPILS6 is an auxin efflux carrier required for maize root morphogenesis"

### Supplemental Figure 1

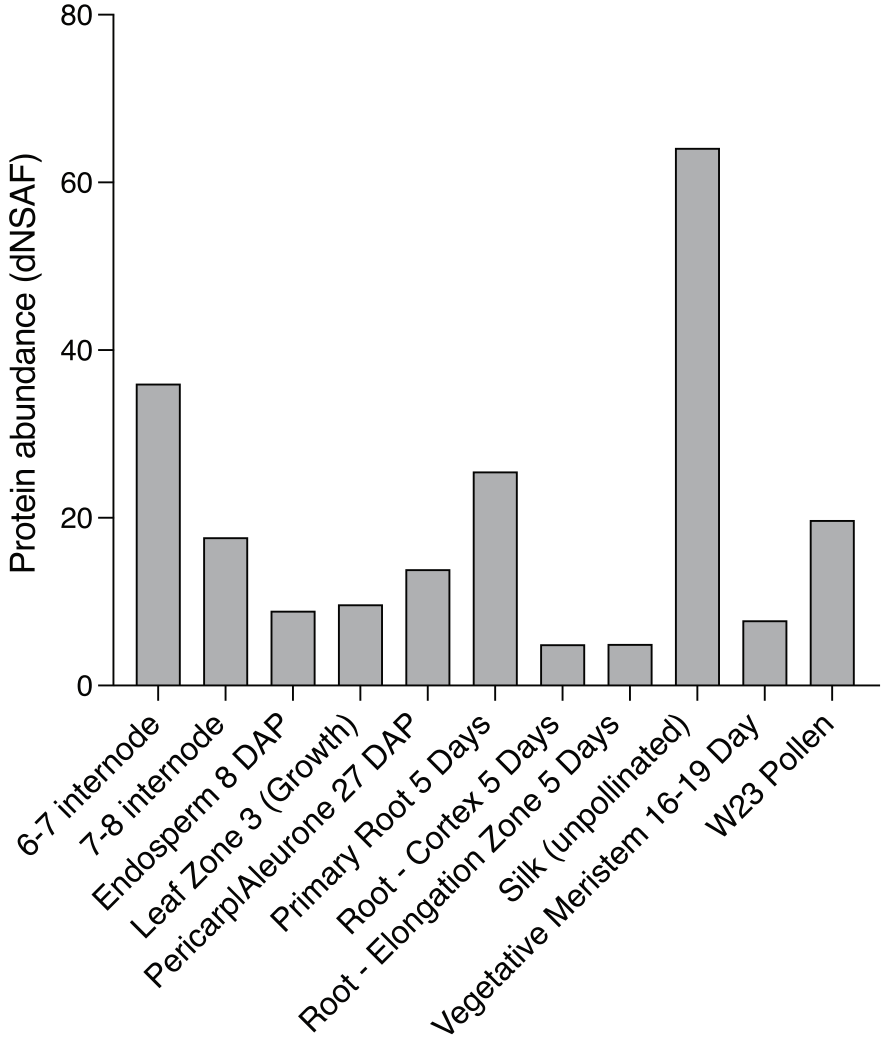

### Supplemental Figure 2

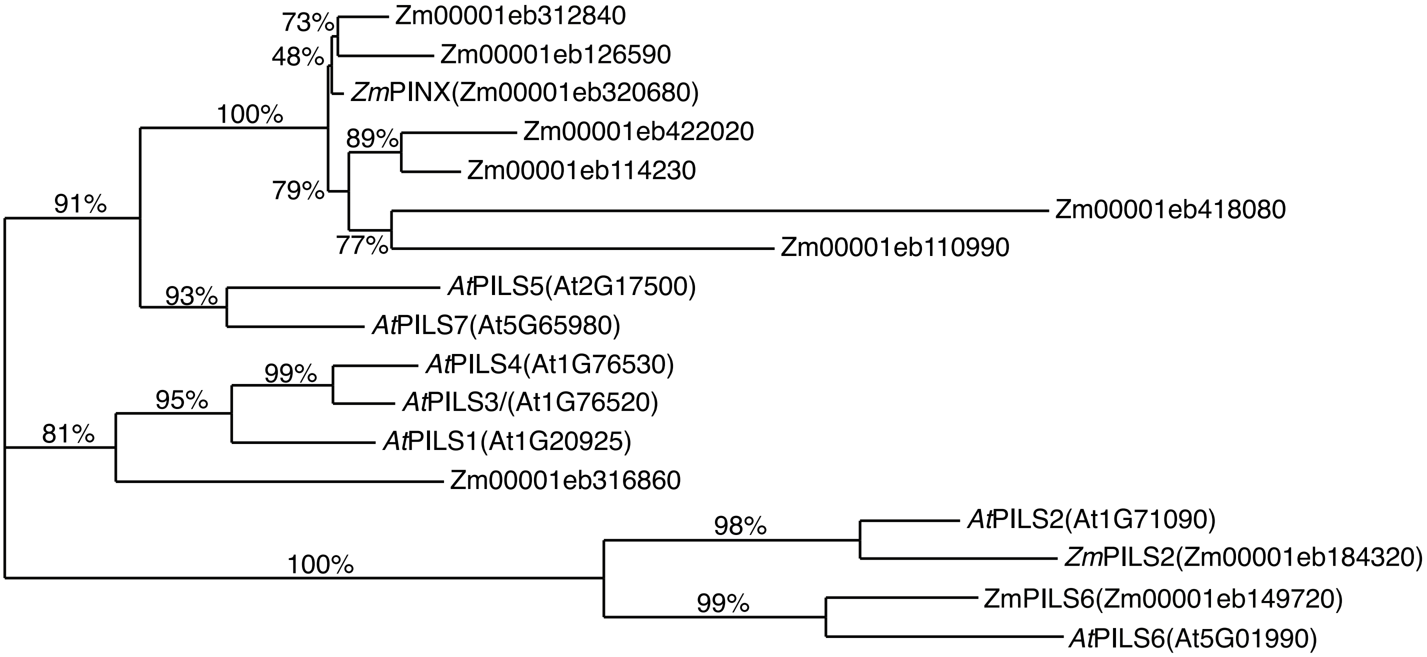

### Supplemental Figure 3

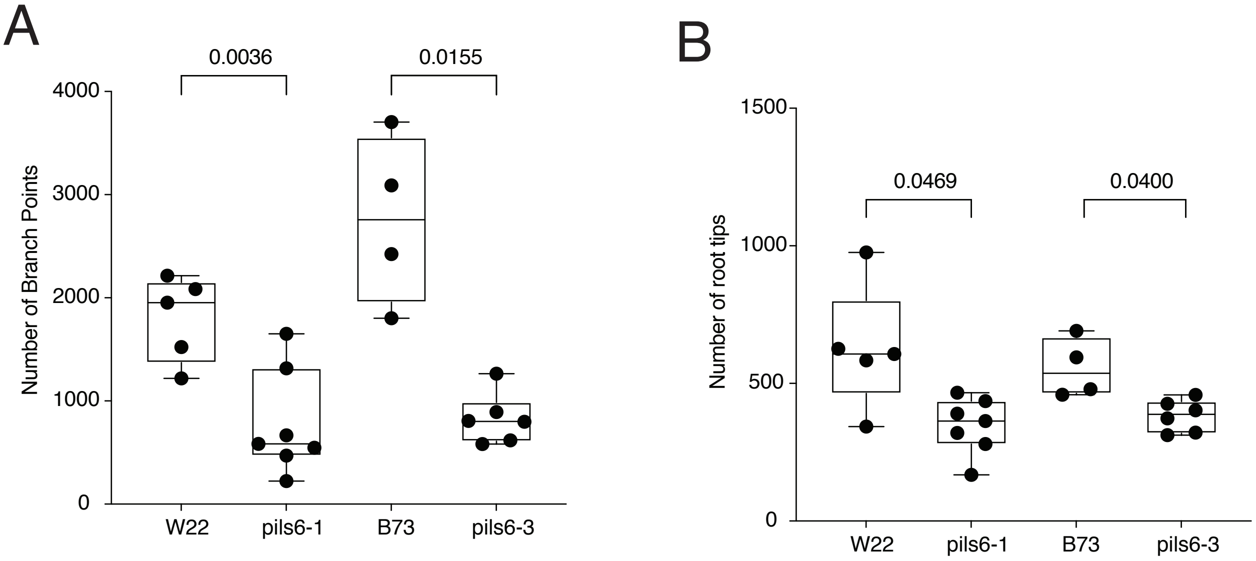

### Supplemental Figure 4

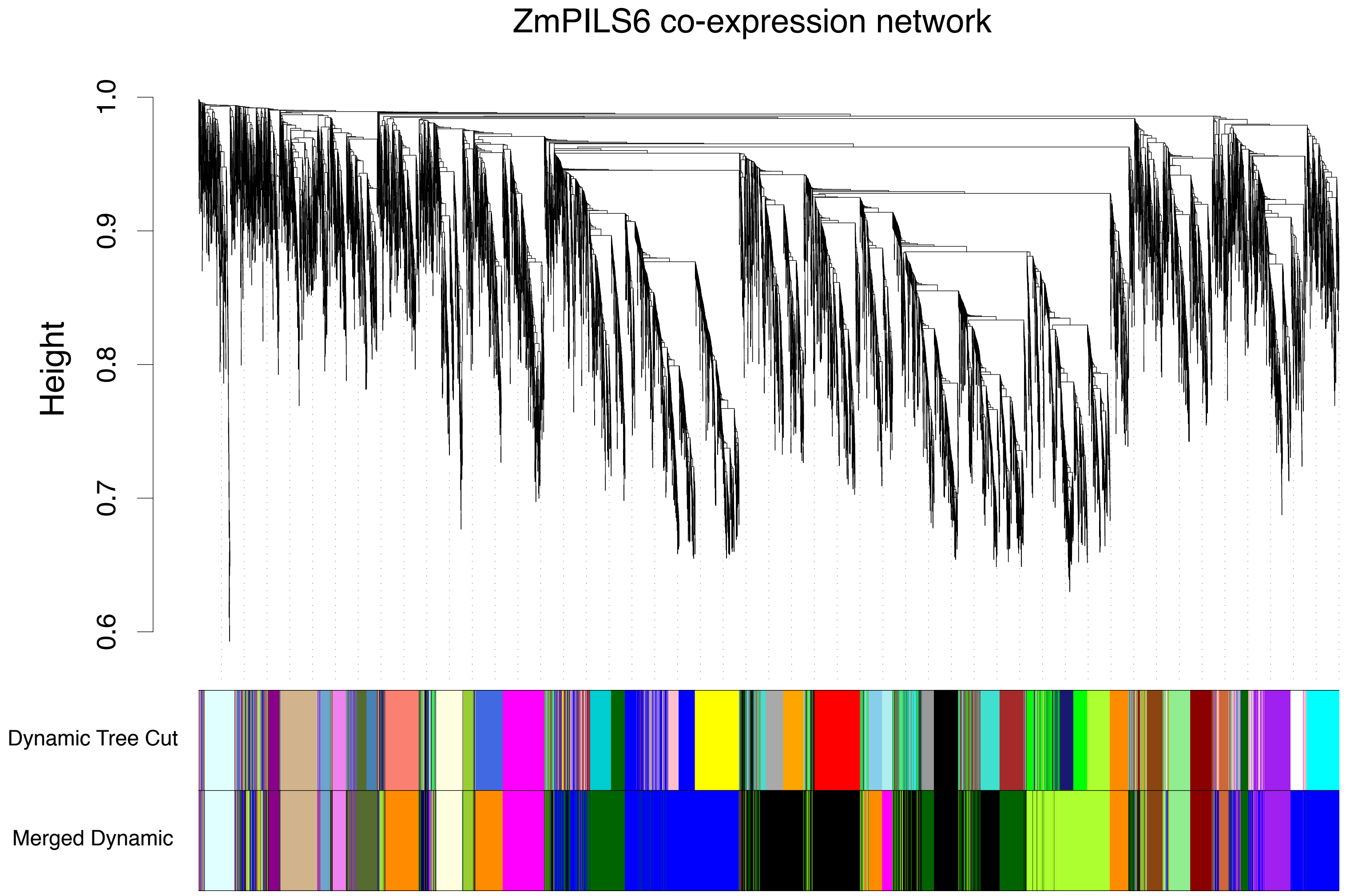
